# Trajectories of atherosclerotic cardiovascular disease risk scores as a predictor for incident chronic kidney disease

**DOI:** 10.1101/2023.06.20.545824

**Authors:** Hye Sun Lee, Hong Il Lim, Tae Ju Moon, So Young Lee, Jun-Hyuk Lee

**Affiliations:** Biostatistics Collaboration Unit, Department of Research Affairs, Yonsei University College of Medicine, Seoul 03277, Republic of Korea; Department of Internal Medicine, Nowon Eulji Medical Center, Eulji University School of Medicine, Seoul 01830, Republic of Korea; Department of Family Medicine, Nowon Eulji Medical Center, Eulji University School of Medicine, Seoul 01830, Republic of Korea; Department of Medicine, Hanyang University Graduate School of Medicine, Seoul 04763, Republic of Korea

**Author notes:** **Corresponding authors’ contact information:** Jun-Hyuk Lee, MD, Assistant Professor, Department of Family Medicine, Nowon Eulji Medical Center, Eulji University School of Medicine, Seoul 01830, Republic of Korea.

**Keywords:** atherosclerotic cardiovascular disease, chronic kidney disease, incidence, prospective cohort

## Abstract

**Background:** The relationship between atherosclerosis and renal function is well established. Atherosclerotic cardiovascular disease (ASCVD) risk scores reflect atherosclerotic burden, which changes over time. We investigated the association between ASCVD risk trajectories and incident chronic kidney disease (CKD) using data from a large community-based Korean cohort with up to 16 years of follow-up.

**Methods:** We analyzed data from 5032 participants without CKD from the baseline survey of the Korean Genome and Epidemiology Study Ansan-Ansung cohort. Participants were categorized into stable or increasing ASCVD risk groups based on the revised ASCVD risk pooled cohort equation over a median period of exposure of 5.8 years. Incident CKD was defined as two consecutive events of an estimated glomerular filtration rate <60 mL/min/1.73 m^2^.

**Results:** During a median 9.9 years of event accrual period, 449 (8.92%) new-onset CKD cases were identified. Multiple Cox proportional regression analyses showed that the hazard ratio (95% confidence interval) for incident CKD in the increasing group, compared to the stable group, was 2.13 (1.74–2.62) in the unadjusted model and 1.35 (1.02–1.78) in the fully-adjusted model. Significant relationships were maintained in subgroups of individuals in their 50s, without diabetes mellitus or hypertension. The prevalence of proteinuria was consistently higher in the increasing group than that in the stable group.

**Conclusions:** An increasing trend in ASCVD risk scores independently predicted adverse renal outcomes. Continuous monitoring of ASCVD risk is not only important for predicting cardiovascular disease but also for predicting CKD.

## Introduction

Atherosclerosis is an important contributor to cardiovascular disease (CVD), including myocardial infarction, ischemic stroke, and peripheral vascular disease.^1, 2^ Atherosclerotic cardiovascular disease (ASCVD) risk scores, such as the Framingham risk score or ASCVD pooled cohort equation, have been developed to estimate an individual’s risk of CVD.^3, 4^ ASCVD pooled cohort equations, in particular, provide race- and sex-specific 10-year ASCVD risk estimates, highlighting the variability in risk factor prevalence, associated risks, and underlying ASCVD event rates among different groups.^3^

There is an inter-relationship between atherosclerosis and renal function.^5, 6^ The pathological process of atherosclerosis can stimulate intrarenal atherogenesis through the mediation of oxidative stress and inflammation, subsequently leading to a decline in renal function.^5^ Concurrently, a decrease in renal function, especially as seen in chronic kidney disease (CKD), can exacerbate the production of uremic toxins, free radicals, and pro-inflammatory cytokines.^6^ Previous research has also suggested that 10-year ASCVD risk scores have predictive power beyond those of conventional cardiovascular risk predictors in determining cardiovascular events in patients with CKD.^7-11^

Given the irreversible nature of CKD, early identification and risk management of high-risk individuals are crucial for preventing its development. Considering the interplay between atherosclerosis and renal function, we hypothesized that the changing trend in ASCVD risk scores might serve as a latent indicator of individuals at a heightened risk of developing CKD. However, to the best of our knowledge, evidence on the relationship between temporal changes in ASCVD risk scores and CKD development is lacking.

If temporal trends in ASCVD risk scores can serve as significant predictors of CKD, they could provide additional information for clinicians to identify individuals at risk of CKD, in addition to those at risk of CVD, allowing for the application of cost-effective early interventions. Therefore, this study aimed to investigate the association between the trajectories of ASCVD risk scores and incident CKD using a large community-based prospective Korean cohort with up to 16 years of follow-up.

## Methods

### Study population

The analysis utilized data from the Ansan-Ansung cohort of the Korean Genome and Epidemiology Study (KoGES), which is a community-based, prospective cohort study conducted by the Korea Centers for Disease Control Agency (KCDA).^12^ In the initial survey conducted in 2001–2002, a total of 10,030 participants aged 40–69 years who had been living in urban (Ansan) and rural (Ansung) areas for at least 6 months were recruited. These participants were followed up biennially for a maximum duration of 16 years, until 2017–2018.

During the baseline survey, data, including anthropometric measurements, personal dietary information, personal medical history, and blood and urine samples, were collected from each participant. The participants were observed from the baseline survey date until the occurrence of the first two consecutive CKD events, the end of the study, or the last date of contact with the participant that provided informative data. The period from the baseline survey to six years was defined as the exposure period, whereas the period from six to 16 years was defined as the event accrual period.

Figure 1 shows a flowchart of the study population selection. Among a total of 10,030 participants at baseline survey of the KoGES_Ansan and Ansung study, we analyzed data from a total of 5032 participants by applying the exclusion criteria as follows:1) Missing data for assessing 10-year ASVD risk scores at baseline (*n* = 136), 2) Prevalent CKD at baseline (*n* = 220), 3) Use of diuretics or steroid medications (*n* = 22), 4) Newly developed CKD during the exposure period (*n* = 209), 5) Never followed up during the exposure period (*n* = 1158), and 6) Never followed up during the event accrual period (*n* = 3253).

**Figure 1.**
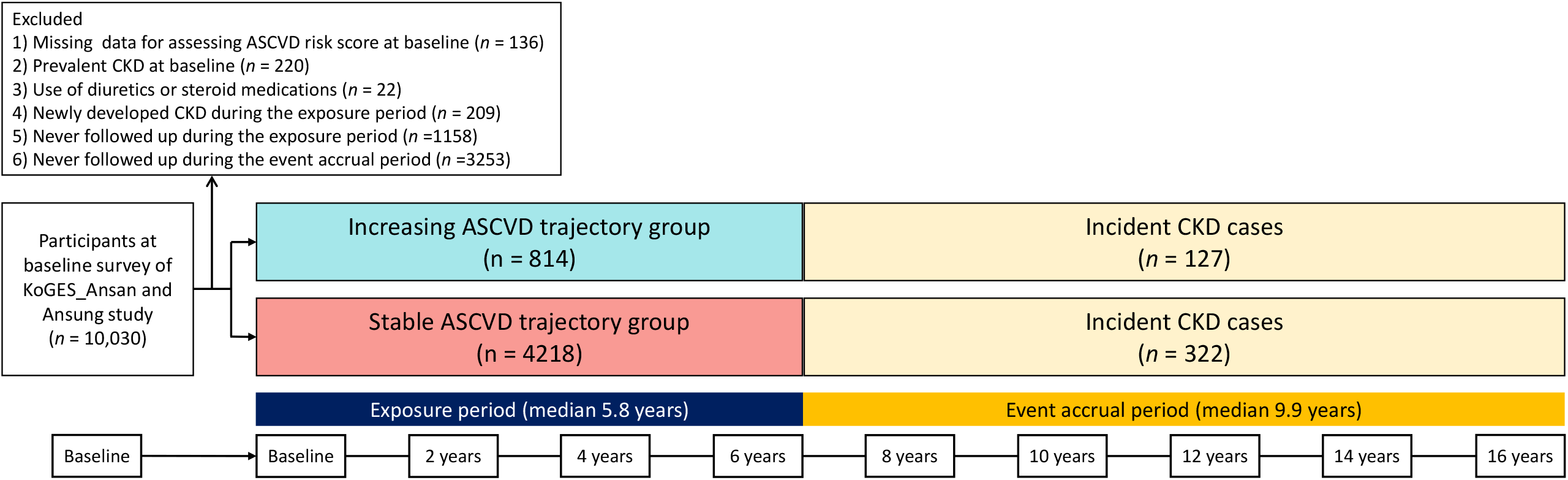
Flowchart of the study population.

The KoGES Ansan-Ansung cohort protocol was reviewed and approved by the Institutional Review Board (IRB) of the KCDA. Written informed consent was obtained from all participants. The study protocol conformed with the ethical guidelines of the 1964 Declaration of Helsinki and its subsequent amendments. This study was approved by the IRB of Nowon Eulji Medical Center (IRB number:2023-02-023).

### Data collection

Height (m) and weight (kg) were measured, and the body mass index (BMI) was calculated as weight divided by height squared (kg/m^2^). Systolic blood pressure (SBP) and diastolic blood pressure (DBP) were calculated as the average of the last two of three measurements. The mean blood pressure (MBP) was calculated. Smoking status was divided into never smoked, ex/ intermittent smokers, and daily smokers. Alcohol consumption status was divided into current drinkers and non-drinkers. Physical activity was measured in metabolic equivalents of task (MET)-hours per day (MET-h/day), based on the International Physical Activity Questionnaire, and was divided into low-intensity (<7.5 MET-h/day), moderate-intensity (7.5–30 MET-hr/day), and high-intensity (>30 MET-h/day) groups. The total energy intake (kcal/day) was calculated using a 103-item food frequency questionnaire. Blood samples were collected from each participant after at least 8-h fasting. Fasting plasma glucose (FPG), serum creatinine, total cholesterol, and C-reactive protein (CRP) levels were also measured. Estimated glomerular filtration rates (eGFR) were calculated using the CKD-Epidemiology Collaboration equation.^13^ Proteinuria was determined based on the results of a urinary dipstick test, with the presence of protein traces of 1+, 2+, 3+, or 4+ considered indicative of proteinuria. The prevalence of proteinuria was calculated to assess the frequency of protein leakage in the study population.

### ASCVD risk trajectories assessment

The main exposure in the current study was the change in ASCVD risk over time. The cardiovascular risk assessment was estimated by using the revised 10-year ASCVD pooled cohort equations.^13^ Using baseline and temporal measurements of the 10-year ASCVD risk scores during the median period of exposure of 5.8 years, we performed trajectory modelling by using R package “lcmm,” which was used for latent variable mixture modelling.^14, 15^ Models with two to five trajectories were constructed by considering the individual as a random effect and time as a fixed effect.^16, 17^ The optimal number of groups of ASCVD risk trajectories was two, based on the Bayesian information criterion and the number of participants in each trajectory (5% or more of the total population), as shown in Supplemental Table 1. Finally, we classified the participants into stable (*n* = 4218) and increasing (*n* = 814) ASCVD risk trajectory groups.

### Renal outcome assessment

The primary outcome of the study was incident CKD, which was defined as the occurrence of two consecutive events with an eGFR of <60 mL/min/1.73m^2^. Participants who did not experience an incident CKD event during the exposure period were followed up from the end of the exposure period until the occurrence of an incident CKD event, the end of the study, or the last date of contact with the participant that provided informative data. The number of new CKD events during the accrual period was also recorded.

### Statistical analysis

Data are presented as mean ± standard deviation for continuous variables and number (percentage, %) for categorical variables. The Student’s t-test was used to compare the differences between the increasing and stable ASCVD risk trajectory groups for continuous variables, whereas the chi-square test was used for categorical variables.

A Kaplan-Meier curve was drawn to compare the cumulative incidence rate of CKD between the stable and increasing ASCVD risk trajectory groups using the log-rank test. Multiple Cox proportional hazard regression analysis was used to estimate the hazard ratio (HR) with a 95% confidence interval (CI) for incident CKD in the increasing ASCVD risk trajectory group compared with the stable ASCVD risk trajectory group. In Model 1, we adjusted for age, BMI, total energy intake, smoking status, alcohol consumption status, and physical activity. In Model 2, we adjusted for the variables used in Model 1 plus MBP, FPG, serum total cholesterol, and CRP levels. In Model 3, we further adjusted for baseline ASCVD risk scores and baseline eGFR. We illustrated a forest plot showing subgroup analyses based on sex, age groups, diabetes mellitus (DM), hypertension (HTN), and high ASCVD risk (≥10%) status, adjusting for variables used in Model 3. The prevalence of proteinuria in the two ASCVD risk trajectory groups at each time point was compared using the chi-squared test. The average changes in eGFR were monitored during the event accrual period, and a comparison was made between the two groups with different ASCVD risk trajectories using a linear mixed model, adjusting for the variables used in Model 3. The interaction between the ASCVD risk trajectory groups and time was analyzed. Sensitivity analysis was also performed for participants who were consistently followed-up during the exposure period.

All statistical analyses were performed using SAS statistical software (version 9.4; SAS Institute Inc., Cary, NC, USA), SPSS statistical software (version 25.0; SPSS Inc., Chicago, IL, USA), and R software (version 4.2.1; R Foundation for Statistical Computing, Vienna, Austria). Statistical significance was set at p < 0.05.

## Results

### Clinical characteristics of the study population based on the different ASCVD risk trajectory groups

Table 1 presents the baseline characteristics of the study population based on ASCVD risk trajectory groups. Compared with the stable ASCVD risk trajectory group, the increasing ASCVD risk trajectory group had a higher proportion of men, individuals aged 50 to 69 years, ex-smokers, intermittent smokers, daily smokers, current drinkers, individuals with high-level physical activity, lower baseline eGFR, higher MBP, FPG, serum total cholesterol level, CRP level, and baseline ASCVD risk scores.

**Table 1.** Baseline characteristics of the study population

| Variables | Total population<br>(*n* = 5032) | ASCVD trajectory groups |  | *p*<sup>*</sup> |
| --- | --- | --- | --- | --- |
|  |  | Stable<br>(*n* = 4218) | Increasing<br>(*n* = 814) |  |
| Male sex, *n* (%) | 2667 (53.0%) | 1776 (42.1%) | 589 (72.4%) | <0.001 |
| Age groups, *n* (%) |  |  |  | <0.001 |
| 40s | 2580 (58.4%) | 2363 (62.5%) | 217 (34.2%) |  |
| 50s | 1434 (32.5%) | 1166 (30.8%) | 268 (42.2%) |  |
| 60s | 403 (9.1%) | 253 (6.7%) | 150 (23.6%) |  |
| BMI, kg/m<sup>2</sup> | 24.6 ± 3.0 | 24.6 ± 3.0 | 24.7 ± 3.1 | 0.244 |
| MBP, mmHg | 95.9 ± 12.8 | 95.2 ± 12.9 | 99.8 ± 11.8 | <0.001 |
| Smoking status, *n* (%) |  |  |  | <0.001 |
| Never smoker | 3165 (62.9%) | 2849 (67.5%) | 316 (38.8%) |  |
| Ex-smoker | 755 (15.0%) | 596 (14.1%) | 159 (19.5%) |  |
| Intermittent smoker | 111 (2.2%) | 81 (1.9%) | 30 (3.7%) |  |
| Daily smoker | 1001 (19.9%) | 692 (16.4%) | 309 (38.0%) |  |
| Current drinker, *n* (%) |  | 1957 (46.6%) | 455 (56.0%) | <0.001 |
| Physical activity, *n* (%) |  |  |  | <0.001 |
| Low | 326 (6.7%) | 273 (6.7%) | 53 (6.8%) |  |
| Moderate | 2953 (60.7%) | 2572 (63.0%) | 381 (48.8%) |  |
| High | 1586 (32.6%) | 1239 (30.3%) | 347 (44.4%) |  |
| Total energy intake, kcal/day | 1970.3 ± 697.7 | 1969.4 ± 699.2 | 1975.2 ± 690.3 | 0.830 |
| Baseline eGFR, mL/min/1.73m<sup>2</sup> | 93.7 ± 12.6 | 94.2 ± 12.6 | 90.7 ± 12.2 | <0.001 |
| FPG, mg/dL | 86.4 ± 19.1 | 85.8 ± 18.5 | 89.9 ± 21.4 | <0.001 |
| Total cholesterol, mg/dL | 190.1 ± 34.3 | 189.4 ± 34.2 | 193.8 ± 34.5 | <0.001 |
| CRP, mg/dL | 0.14 [0.06;0.23] | 0.13 [0.06;0.23] | 0.16 [0.08;0.27] | <0.001 |
| Baseline ASCVD risk, % | 2.3 [0.7;5.5] | 1.8 [0.5; 4.4] | 6.2 [3.4;10.7] | <0.001 |

\*p-values were derived from Student's t-test or Mann-Whitney test for continuous variables and chi-square test for categorical variables to compare differences in variables between groups. A p-value of less than 0.05 was considered statistically significant.
Abbreviations: ASCVD, atherosclerotic cardiovascular disease; BMI, body mass index; MBP, mean blood pressure; eGFR, estimated glomerular filtration rate; FPG, fasting plasma glucose; CRP, C-reactive protein.

### Association of ASCVD risk trajectory groups with incident CKD during the event accrual period

During the median 9.9 years of the event accrual period, a total of 449 (8.92%) newly developed CKD cases occurred. The incidence rate of CKD per 1000 person-years was 9.35.

Figure 3 shows the Kaplan-Meier curves for the cumulative incidence rate of CKD during the event accrual period according to the two different ASCVD risk trajectory groups. The increasing ASVD risk trajectory group showed a persistently higher cumulative incidence of CKD than the stable ASVD risk trajectory group (*p* for log-rank test <0.001). Table 2 presents the Cox proportional hazard regression analysis for incident CKD in the increasing ASCVD risk trajectory group compared with the stable ASCVD risk trajectory group. In the unadjusted model, the HR (95% CI) for incident CKD in the increasing ASCVD risk trajectory group compared with that in the stable group, was 2.13 (1.74– 2.62). Significant relationships remained in the adjusted model. The adjusted HR (95% CI) for incident CKD in the increasing ASCVD risk trajectory group, compared to stable group, was 1.39 (1.06–1.82) in Model 1, 1.35 (1.02–1.77) in Model 2, and 1.35 (1.02–1.78) in Model 3. Subgroup analysis revealed a significant association between ASCVD risk trajectories and incident CKD in individuals in their 50s without DM and HTN (Supplementary Figure 2).

**Table 2.** Cox proportional hazard regression analysis showing the relationship of ASCVD risk trajectories with incident chronic kidney disease

|  | Stable ASCVD risk | Increasing ASCVD risk |  |  |
| --- | --- | --- | --- | --- |
| Total cases, n | 4218 | 814 |  |  |
| Incident CKD cases, n | 322 | 127 |  |  |
| Follow-up time, person-year | 40492.7 | 7536.8 |  |  |
| Incident rate per 1000 person-year | 7.95 | 16.85 |  |  |
|  |  | HR | 95% CI | *p* |
| Unadjusted | 1 (reference) | 2.13 | 1.74–2.62 | <0.001 |
| Model 1 | 1 (reference) | 1.39 | 1.06–1.82 | 0.016 |
| Model 2 | 1 (reference) | 1.35 | 1.02–1.77 | 0.033 |
| Model 3 | 1 (reference) | 1.35 | 1.02–1.78 | 0.033 |

Model 1: Adjusted for sex, age groups, BMI, total energy intake, smoking status, alcohol drinking status, and physical activity.
Model 2: Adjusted for variables used in Model 1 plus MBP, FPG, serum total cholesterol, and CRP levels.
Model 3: Adjusted for variables used in Model 2 plus baseline ASCVD risk scores and baseline eGFR.
Abbreviations: ASCVD, atherosclerotic cardiovascular disease; CKD, chronic kidney disease; BMI, body mass index; MBP, mean blood pressure; FPG, fasting plasma glucose; CRP, C-reactive protein; eGFR, estimated glomerular filtration rate; HR, hazard ratio; CI, confidence interval.

**Figure 2.**
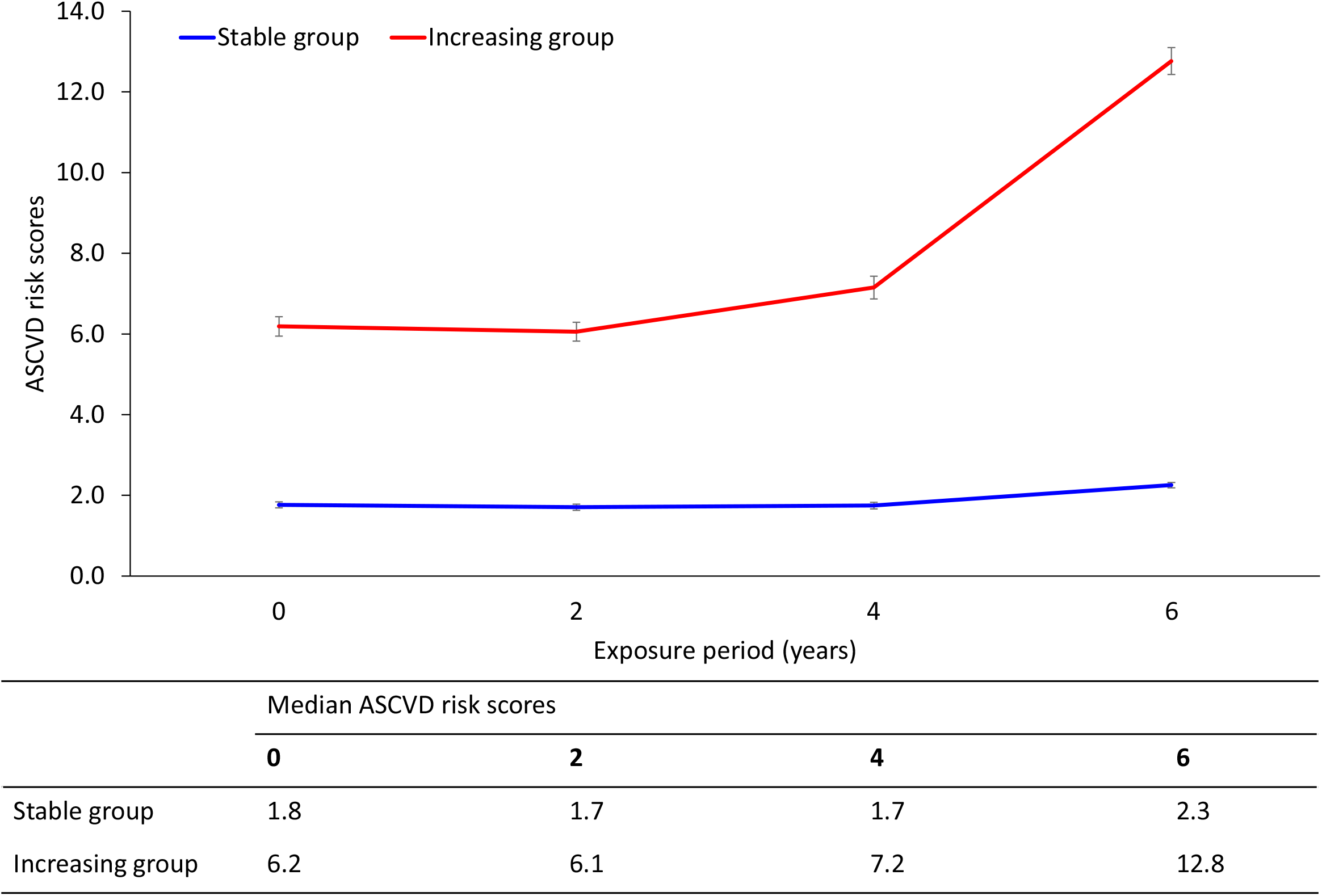
Trajectory modeling with the revised ASCVD risk pooled cohort equation using the latent class linear mixed model. Red line: increasing ASCVD risk trajectory group, blue line: stable ASCVD risk trajectory group. Abbreviation: ASCVD, atherosclerotic cardiovascular disease.

**Figure 3.**
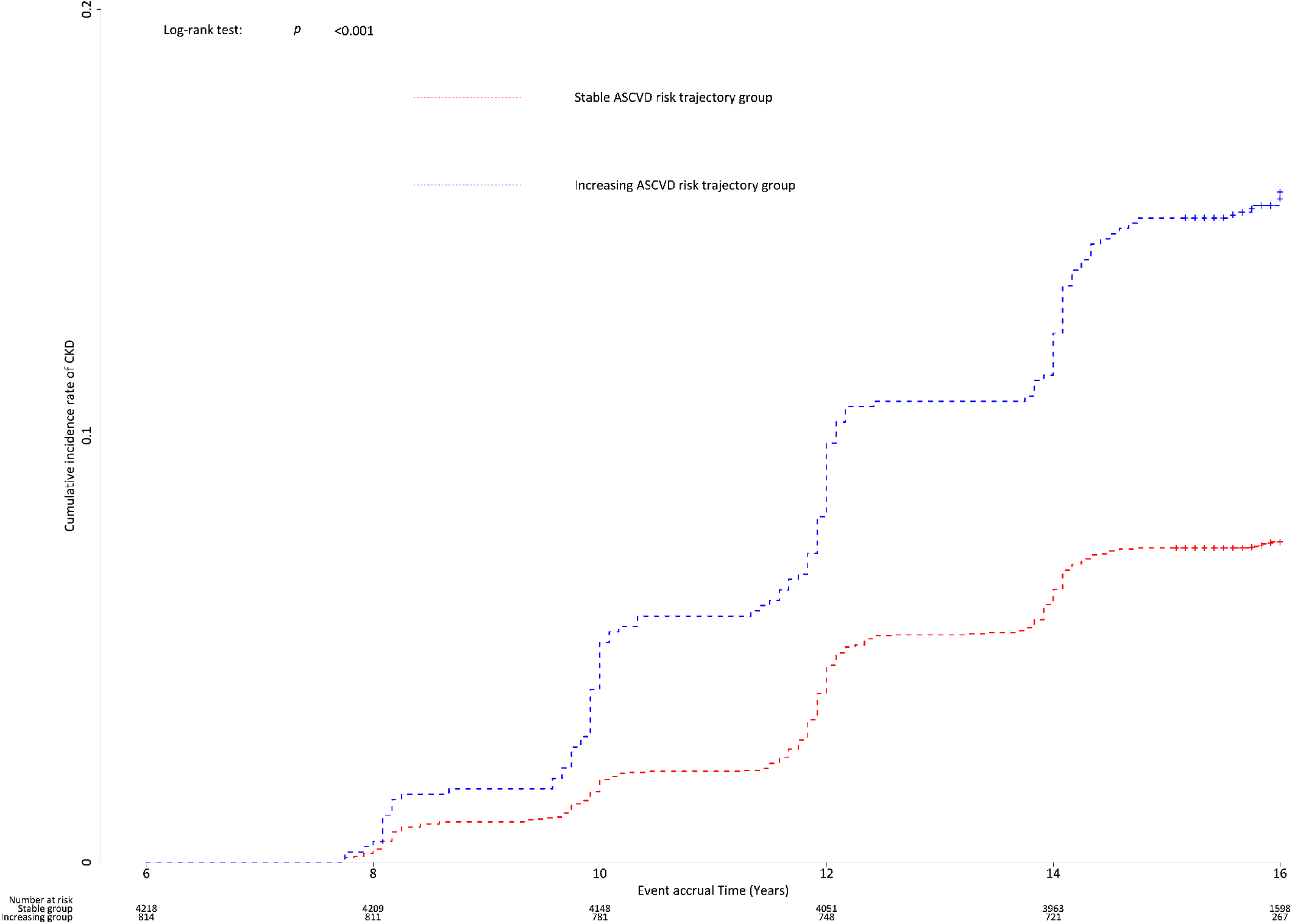
Kaplan-Meier curves showing the cumulative incidence rates of CKD based on the ASCVD risk trajectory groups. Abbreviations: CKD, chronic kidney disease; ASCVD, atherosclerotic cardiovascular disease

### Prevalence of proteinuria between the increasing and stable ASCVD risk trajectory groups

Table 3 presents the prevalence of proteinuria based on the ASCVD risk trajectory groups during the event accrual period. The prevalence of proteinuria was consistently and significantly higher in the increasing ASCVD risk trajectory group than that in the stable group during the event accrual period.

**Table 3.** The prevalence of proteinuria according to the ASCVD risk trajectory groups.

| Prevalence of proteinuria (%) | ASCVD risk trajectory groups |  | *p* |
| --- | --- | --- | --- |
|  | Stable | Increasing |  |
| At 8 years | 1.90 | 4.02 | 0.004 |
| At 10 years | 1.75 | 3.46 | 0.015 |
| At 12 years | 3.40 | 5.71 | 0.013 |
| At 14 years | 4.03 | 8.31 | <0.001 |
| At 16 years | 6.73 | 9.94 | 0.005 |

Abbreviations: ASCVD, atherosclerotic cardiovascular disease.

### Longitudinal changes in eGFR according to the ASCVD risk trajectory groups

In the linear mixed model, the increasing ASCVD risk trajectory group had a consistently lower eGFR than the stable ASVD risk trajectory group (Supplementary Figure 1). The significant interaction between group and time indicated that the rate of change in eGFR over time differed between the groups. In the post hoc analysis, there were significant differences in the GFR between the stable and increasing ASCVD risk trajectory groups during all event accrual periods.

### Sensitivity analysis with participants who were consistently followed up during the exposure period

During the exposure period, 4634 participants were consistently followed up. Among these, 411 cases (8.87%) of newly developed CKD were identified. Supplementary Table 2 demonstrates a higher risk of incident CKD in the increasing compared to the stable ASCVD risk trajectory group, with fully-adjusted HR of 1.36 and 95% CI of 1.03–1.81. The prevalence of proteinuria was consistently higher in the increasing ASCVD risk trajectory group than that in the stable group (Supplementary Table 3).

## Discussion

To the best of our knowledge, this is the first study to investigate the association between ASCVD risk trajectory and incident CKD. Previous studies have concentrated mainly on evaluating ASCVD risk in individuals with pre-existing CKD.^18-20^ However, the identification of individuals at a higher risk of CKD through early detection and implementation of early interventions can be a potentially cost-effective approach to reducing the burden of CKD. Considering the interrelationship between ASCVD and CKD,^5, 6^ we hypothesized that the trajectories of ASCVD risk scores could predict the risk of development of CKD.

In this prospective cohort study of 5032 Korean adults, the ASCVD risk trajectory group had a 2.13 times higher risk of incident CVD than the stable group, regardless of the ASCVD risk at baseline. Moreover, the increasing group consistently showed a higher prevalence of proteinuria than did the stable group during the event accrual period. These findings highlight the importance of considering the ASCVD risk trajectories when predicting proteinuria. Mean eGFR was consistently lower in the increasing than in the stable ASCVD risk trajectory group. At the end of the follow-up period, both groups showed a rapid increase in the mean eGFR. This observation suggests that a sufficient number of CKD cases may have occurred during the study, leading to only individuals with healthy renal function remaining in the analysis.

Ren et al.^8^ reported an association between the ASCVD risk and biopsy-proven CKD in patients with type 2 DM. Among 218 patients with type 2 DM and biopsy-proven CKD, individuals with higher ASCVD risk scores showed lower eGFR and more severe renal interstitial inflammation. They also suggested that an ASVD risk of 14.1% was an independent indicator of renal dysfunction. In the current study, similar to a previous study, the increasing ASCVD risk trajectory group had higher baseline ASCVD risk scores and a lower eGFR than the stable ASCVD risk trajectory group. At the end of the exposure period, the median ASCVD risk score in the increasing ASVD risk trajectory group was 12.8, which was lower than the risk score suggested previously. As we investigated the risk of development of CKD in the future using participants without CKD, the ASCVD risk score for predicting renal dysfunction may be lower than that suggested previously.

The risk of incident CKD was associated with ASCVD risk trajectories in the subgroups of patients in their 50s, those without DM, and those without HTN. The incidence of CVD is approximately 40% in the age group of 40–59 years and 75% in the age group of 60–79 years.^21^ In a study that tracked middle-aged men over a period of 40 years, the influence of traditional CVD risk factors declined with age and only low-density lipoprotein cholesterol consistently affected CVD risk, while BMI and FPG had an impact on heart failure, regardless of age.^22^ Given that age in itself is a significant risk factor for both CVD and CKD, the lack of a significant correlation between the increase in ASCVD risk score in individuals in their 60s and the risk of incident CKD in our study could suggest that age may have played a vital role in the development of CKD. In contrast, the incidence rates of CKD per 100,000 population years were 132 for mild CKD, 33 for moderate CKD, and 15 for severe CKD in individuals aged 20 to 64 years.^23^ Participants from the current study aged 40–49 years may not have been able to demonstrate the impact of changes in the ASCVD risk score on the incidence of CKD due to low CKD rates even after a period of 16 years, given that they remained below the age of 65 at completion of the study. In contrast, we hypothesized that, in individuals aged 50–59 years, we were able to establish a significant correlation between alterations in ASCVD risk scores and the incidence of CKD. This is likely related to their reaching an increased risk of CKD during the substantial 16-year follow-up period, whereas the effects of CVD risk factors did not diminish with age. In patients with DM, those individuals with DM at baseline could have already developed CKD to a sufficient extent during the exposure period, given that it takes approximately 7–10 years for CKD to develop. ^24^ The rapid decline in renal function was observed particularly in patients with DM with high SBP.^25^ Similarly, in patients with HTN, an analysis using data from a total of 43,305 patients with HTN from the United States (US) revealed that time-varying SBP is a factor for a rapid decline in eGFR, with every 10 mmHg increase in SBP leading to a 0.2 mL/min/1.73 m^2^ decline in eGFR.^26^ Furthermore, an elevated risk for the onset of CKD was observed in patients with HTN who maintained a SBP of 120 mmHg or above.^26^ Similarly, in a Korean study, SBP ≥130 mmHg and DBP ≥90 mmHg were related to an increased risk of incident CKD.^27^ As the formula for calculating ASCVD risk scores includes SBP, individuals who showed an increasing trend in ASCVD risk scores may have been excluded due to the early onset of CKD. This could have attenuated the association between changes in ASCVD and the incidence of CKD during the event accrual period in patients with DM or HTN.

We suggest several explanations for these findings, as well as providing the exact underlying mechanism for the significant relationship between ASCVD risk trajectory groups and incident CKD. First, both ASCVD and CKD share several common risk factors such as HTN, DM, and dyslipidemia.^28^ It is possible that individuals in the group with increased ASCVD risk experience these comorbidities, leading to an increased risk of CKD. At the molecular level, a recent study suggested that microRNAs (miRNAs) mediate homeostasis of the vascular wall and are involved in vasculature pathologies.^29^ Certain miRNAs, such as miR-155 and miR-223, play a role in the imbalance of calcium and phosphate seen in the vessels. This imbalance leads to the loss of these minerals from the bones and their accumulation in the vascular wall, contributing to vascular calcification, a common complication of CKD. In addition, miR-145 is upregulated in the plasma of patients with CVD and, along with miR-143, is involved in angiogenesis and EC proliferation. Finally, miR-21 is upregulated in endothelial cells from atheroma plaques and progenitor cells from patients with coronary artery disease and is associated with the physiological proliferation of vascular smooth muscle cells and kidney fibrosis.^25^

Second, an increase in ASCVD risk could reflect a more severe or active atherosclerotic disease process, thus leading to higher susceptibility to renal damage and subsequent CKD.^5, 30^ Systemic inflammation plays a crucial role in the development and progression of atherosclerosis throughout the vascular system, including the blood vessels in the kidneys.^5^ Endothelial dysfunction limits vascular relaxation and causes infiltration of inflammatory cells.^30^ Cigarette smoking, a component of the formula for the revised ASCVD pooled equation, can exacerbate endothelial dysfunction and facilitate the infiltration of inflammatory cells into the vascular wall, thus promoting the formation of atherosclerotic plaques. In our study, we observed a higher proportion of daily smokers and elevated CRP levels in the group with increasing ASCVD risk trajectory than that in the stable group. These findings support the hypothesis that the increasing ASCVD risk trajectory group may have a higher risk of renal arterial atherosclerosis. This, in turn, could contribute to the observed association between an increased ASCVD risk trajectory and CKD incidence. Additionally, other lifestyle factors such as physical inactivity and alcohol consumption may contribute to the progression of ASCVD, which may lead to renal damage over time.

This study had several limitations. First, the exclusion of individuals with new-onset CKD during the exposure period may have limited the ability of our trajectory modeling to fully reflect the risk of CKD onset. However, we believe that we have partially addressed the limitation of excluding early-onset CKD cases by obtaining a sufficient number of new-onset CKD cases over a 10-year event-accrual period. Second, the risk of CKD may have been underestimated in the increasing ASCVD risk trajectory group. Considering that the mean baseline ASCVD risk score was higher in the increasing than in the stable ASCVD risk trajectory group, individuals with an increasing ASCVD risk trajectory may experience CVD mortality before developing CKD. Third, our results may not be applicable to other ethnicities because the KoGES consists only of Koreans. Finally, we focused solely on changes in ASCVD risk scores over time and did not account for the dynamic effects of other important variables such as BMI, blood pressure, plasma glucose level, lipid profiles, and lifestyle factors. Further studies incorporating these factors and their time-dependent effects will provide valuable insights into the development and progression of CKD.

In conclusion, our study provides evidence that an increasing trend in ASCVD risk scores serves as an independent predictor of adverse renal outcomes such as new-onset CKD and the presence of proteinuria. Our findings underscore the importance of continuous monitoring and effective control of ASCVD risk factors, not only for cardiovascular health but also for the prevention and management of CKD. Further research is needed to validate these findings and elucidate the precise mechanisms underlying this association.

## Non-standard abbreviations and acronyms

ASCVD: atherosclerotic cardiovascular disease
CKD: chronic kidney disease
CRP: C-reactive protein
CVD: cardiovascular disease
DM: diabetes mellitus
eGFR: estimated glomerular filtration rate
HTN: hypertension
FPG: fasting plasma glucose
IRB: institutional review board
KCDA: Korea Centers for Disease Control Agency
KoGES: Korean Genome and Epidemiology Study
MET: metabolic equivalents of task

## Acknowledgments

Data in this study were from the Korean Genome and Epidemiology Study (KoGES; 6635-302), National Institute of Health, Korea Disease Control and Prevention Agency, Republic of Korea.

## Data availability

The dataset used in this study can be provided after a KCDA review and evaluation of the research plan (http://www.nih.go.kr/ko/main/contents.do?menuNo=300563).

## Author Contributions

Hye Sun Lee: study concept and design; methodology; acquisition, analysis, and interpretation of data; drafting the manuscript; Hong Il Lim: interpretation of data; Tae Ju Moon: interpretation of data; So Young Lee: interpretation of data; methodology; Jun-Hyuk Lee: study concept and design; methodology; interpretation of data; supervision; revising the manuscript; All authors approved he final manuscript.

## Ethical approval

The KoGES_Ansan_Ansung cohort protocol was reviewed and approved by the Institutional Review Board (IRB) of the KCDA. All the participants read and signed a written informed consent form. The study protocol conformed to the ethical guidelines of the 1964 Declaration of Helsinki and its later amendments. This study was approved by the IRB of the Nowon Eulji Medical Center (IRB number: 2023-02-023).

## Novelty and Significance

What is known?

- The relationship between atherosclerosis and renal function is well established.
- Atherosclerotic cardiovascular disease (ASCVD) risk scores reflect atherosclerotic burden, which changes over time.

What new information does this study contribute?

- In a group of patients with increasing ASCVD risk scores, there was a 2.13-fold increase in incident chronic kidney disease
- The prevalence of proteinuria was higher in patients with increasing ASCVD risk scores over time.
- An increasing trend in ASCVD risk scores serves as an independent predictor of adverse renal outcomes.

While the association between atherosclerosis and renal function is well known, the atherosclerosis cardiovascular disease (ASCVD) risk scores have been applied only to cardiac disease. We thus aimed to assess whether these scores might be used to predict chronic kidney disease (CKD). Using data from a large community-based Korean cohort with up to 16-year follow-up, our study showed evidence that an increasing trend in ASCVD risk scores can serve as an independent predictor of adverse renal outcomes such as new-onset CKD and the presence of proteinuria. To our knowledge, this is new information which has not been reported previously. In subgroup analysis, the predictive value of ASCVD risk scores applied to patients in their 50s, and those without diabetes mellitus or hypertension. Factors such as age and comorbidities might be independent risk factors for CKD, hence these results. Our findings underscore the importance of continuous monitoring and effective control of ASCVD risk factors, not only for cardiovascular health but also for the prevention and management of CKD. Further research is needed to validate these findings and elucidate the precise mechanisms involved.

